# Six-legged-bound: a newly-described insect gait

**DOI:** 10.1101/2024.11.22.624955

**Authors:** A. Amir, O. Yuval, A. Ayali

## Abstract

Locomotor behavior is a hallmark of animal biology and ecology. Mole crickets constitute a unique group of subterranean insects that present an exceptional model for locomotion-related research. In response to an aversive stimulus from the front, the mole cricket will consistently adopt a unique backwards gait that we have termed backward-bound. This never before reported six-legged gait comprises a cyclic alternation between the middle and hind-leg pairs with left-right in-phase synchronization, while the front legs display noisy and less consistent phase dynamics. This exceptional gait is transient and is usually expressed for up to a dozen cycles. It is employed to distance the animal quickly away from danger. A gait that can be characterized as “forward-bound” is also displayed by the mole cricket, albeit for a much shorter duration (up to two cycles).

## Introduction

Characterizing the locomotor behavior of animals is essential to any study of their biology and ecology (e.g., Irschick and Garland, 2001; Biewener and Patek, 2018). Locomotor behavior constitutes the way in which the animal navigates its environment, i.e., the trajectory of the body within the environment and its interaction with the substrate; as well as the controlled and coordinated way in which the animal moves its body and appendages against each other.

These different aspects of locomotion are of special interest in the case of the mole cricket (Orthoptera: Gryllotalpidae). Mole crickets are subterranean insects and among the few families of insects known to possess fossorial (digging) front legs (Fig. 1A). The enlarged dactyl claws on the insect’s tibia, common to all species of mole crickets, enable their unique soil-dwelling, subterranean life history. The specialized hind legs (Fig. 1A), in addition to serving as jumping legs (as in many other orthopteran insects), provide most of the thrust needed for pushing the animal forward when digging underground (Kidd, 1825). Other modifications related to the mole crickets’ life in an underground network of tunnels and burrows are their cylindrical, sclerotized body, and a pointed head (e.g. Kidd 1825; Zhang et al., 2008, 2019; Fig 1). Interestingly, these unique morphological modifications have already been distinguished in mole cricket specimens preserved in mid-Cretaceous Burmese amber (Wang et al., 2020; Xu et al., 2020).

**Figure 1:**
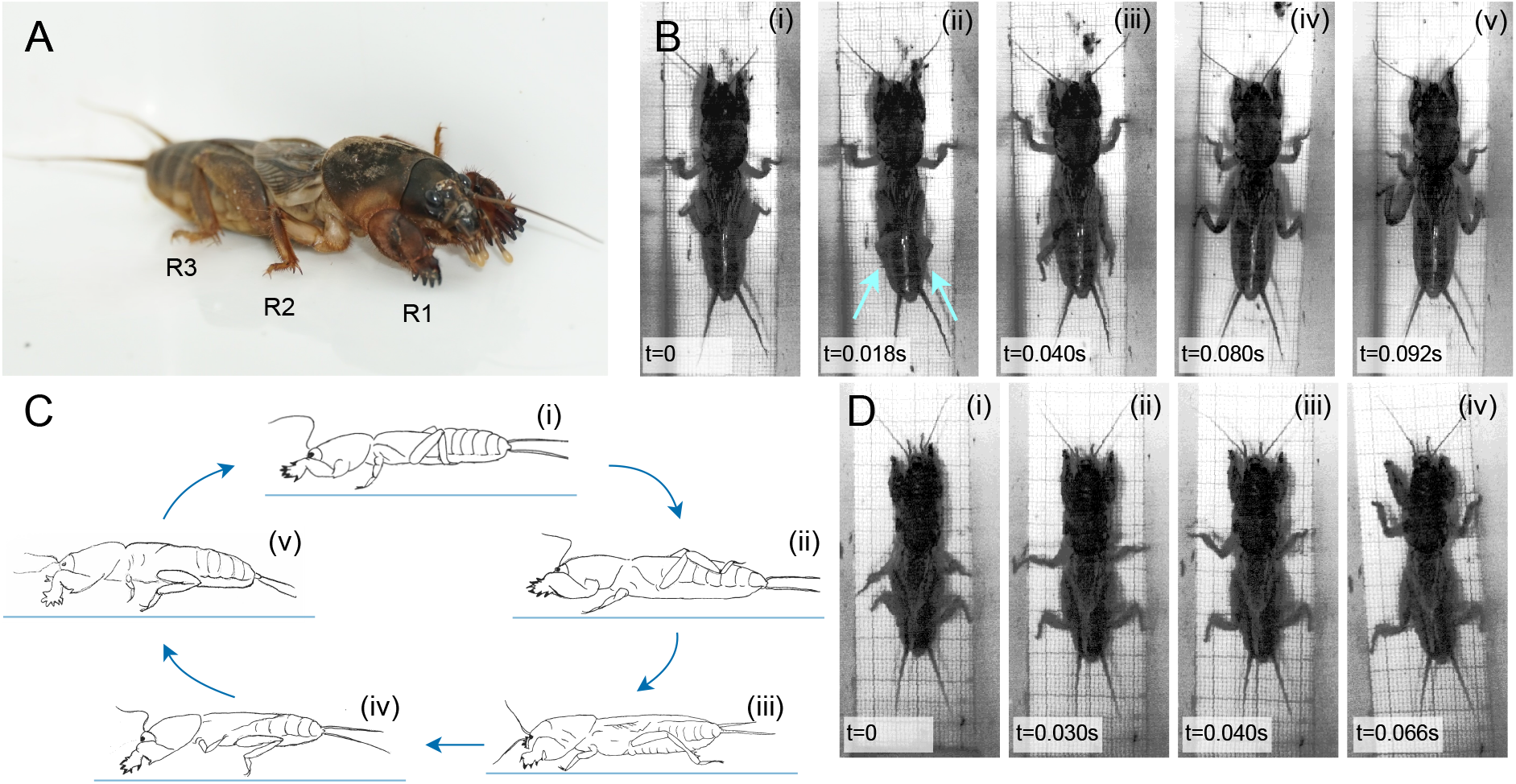
**A**. The mole cricket presents extreme morphological adaptations to its subterranean life. **B**. The backward-bound gait: Sample images extracted from a video used in the presented analysis, highlighting the left-right legs’ symmetry. Arrows depicts hind-leg tips. **C**. An artist’s side view (based on a video) of the backward-bound gait, demonstrating the leg and body trajectories. D. Four main stages in the forward-bound gait (*cf*. B)

Six-legged locomotion is exceptionally effective, making insects (together with their other traits) one of the most successful groups of organisms. One reason for this preeminence is the insects’ remarkable ability for dynamic stability: Insects can rapidly generate adaptable movement in changing environments, employing multi-level adaptations, including adaptive control mechanisms (Graham, 1985, Ritzmann and Büschges, 2007; Ayali et al., 2015a). The dominant and ubiquitously observed insect gait is the double-tripod gait, characterized by distinctive phasing among the legs in terms of time and position: the front and rear legs on one side of the body move in phase with the middle leg on the other side and in anti-phase with their contralateral partners, resulting in two alternating tripods. The double-tripod gait is considered extremely stable and is assumed to be partly responsible for the outstanding fast locomotion of some insects and their ability to negotiate different terrains.

One important aspect of dynamic and adaptable locomotion is manifested in movement-related decision-making, i.e., in selecting the gait most appropriate to the context. For example, in many animals, rapid, intense locomotion is often seen as part of their aversive response to threatening stimuli (Card, 2012; Branco and Redgrave 2020). This type of behavior is mostly transient, and is characterized by sudden, explosive movements allowing the animal to quickly distance itself from the potential danger. Its complex and challenging environment makes the mole cricket a good model system also for the study of distinct, context-specific gait selection.

Here we present a never before reported insect gait that we term “backward-bound”, demonstrated by the mole cricket in response to aversive stimuli. This unique gait is dominated by cycles of alternation between the mid-leg and hind-leg pairs (hind legs leading during backward locomotion), with intra-pair in-phase synchronization. Remarkably, such a temporal sequence has not been observed in any other mode of six-legged locomotion.

## Materials and Methods

### Animal maintenance and experimental procedure

Mole crickets (*Gryllotalpa tali*) were collected from the countryside around Tel Aviv, Israel, at various life stages. Insects were maintained in darkness, at room temperature (20-25°C), each in a separate 750 ml glass jar filled with autoclaved plant soil, and fed with flour beetle larvae, grass roots, and thin slices of carrot. A moist environment was maintained by sprinkling each jar with 100 ml of water every three days. Adult males and females that were acclimated to the laboratory conditions for at least 3 months were used for the experiments.

The imaging setup comprised a 30Lx2.5Wx10H cm tunnel, with a 10×10×10 cm acclimation chamber at either end. A millimeter graph paper was glued to the floor of the tunnel in order to extract a scaling factor to convert from pixels to real length units. The temperature during experiments was maintained at 24±1°C. Video sequences were captured from ca. 20 cm above the setup, using a digital high-speed camera (500 fps; MIKROTRON motion-Blitz cube4MGE-CM4, Germany), fitted with a VS-H1218-IRC/11 lens (Vital Vision Technology Pte Ltd, Singapore). The lighting used to illuminate the setup was covered with red cellophane to avoid glare and reduce stress to the insects.

Insects were introduced individually into one of the acclimation chambers, and their locomotion was recorded for up to 10 minutes while they moved freely (spontaneously) in the tunnel section. In order to induce the bound gait, mild aversive stimuli were applied to the head or the tip of the abdomen in the form of a gentle push with a piece of sponge on a wooden stick. Data for the backward-bound gait comprised 20 individuals, including 10 males and 10 females. Data for the forward-bound gait comprised 8 individuals, including 4 males and 4 females.

### Video analysis and extraction of locomotion parameters

To extract kinematic parameters from the recorded data, key points on the animal’s body were tracked in all videos. These comprised leg tips (tip of the tarsus), leg bases (thorax-coxa junction), and the thorax-head (TH) and thorax-abdomen (TA) junctions (used to define the main body axis). To this end, frames were selected from videos of different gaits, body poses and animals, and the above noted key points were manually labeled in order to train a ResNet50 model. The trained model was then applied to the entire video dataset to detect those points in all frames (n=5036). 230 frames (4.6%) had to be manually corrected, mostly due to cases of self-occlusion or soil particles that were attached to the body. Labeling, training, and tracking were conducted using DeepLabCut (a software package for animal pose estimation. Created by the A. and M.W. Mathis Labs Mathis et al., 2018).

The position of the leg tip of leg i was defined as the shortest distance of the tip of the tarsus from the secondary body axis (perpendicular to the main body axis, with its origin at the TA point). Leg tip position over time, d_i_(t), was used to calculate swing-stance matrices and inter-leg phases. Using the maxima and minima in this signal for each leg, forward stances were defined as the ranges from the anterior-most to the posterior-most leg tip positions, while swing was defined as the rest of the signal. Backward stances were defined in the opposite way (i.e., from the posterior-most to the anterior-most leg tip positions). The inter-leg phase of leg i and j was calculated by computing the cross-correlation between d_i_(t) and d_j_(t).

Leg angle was defined as the angle between the vector connecting the leg base to its tip, and the main body axis. To obtain the average leg angle, the circular mean was first used to average leg angles over time, and then across videos. Similarly, the circular standard deviation was used to obtain the standard deviation of leg angles over time, and then averaged across videos using a regular mean. Statistics for the mean leg angle were calculated using the Kuiper’s test for circular data. Statistics for the standard deviation were calculated using the two-sample t-test.

## Results

Mole crickets in our experimental set-up demonstrated mostly forward walking with occasional backward locomotion, both while showing the characteristic double-tripod gait (see also Zhang et al., 2011; 2019). Upon experiencing an aversive stimulus from the front (in the shape of an approaching decoy comprising a piece of sponge on a wooden stick), while standing or in the midst of a forward locomotion bout, the insect swiftly switched to moving rapidly backwards, while adopting a unique mode of leg coordination that we have termed “backward-bound” (Fig. 1B, C).

Utilizing video tracking and machine learning we analyzed the details of the legs’ movements during backward-bound, as well as the distinct phasing between the legs (Fig. 2A-D, N=20; 10 males and 10 females). This unique gait is characterized by left-right in-phase synchronization between leg pairs, with the two dominant leg pairs – the middle and the hind legs – consistently alternating (Fig. 2B, C, video S1): i.e., while the middle legs are in the stance (pushing against the ground), the hind legs are in swing (raised and moved far back, Figs. 1, 2C,D). Conversely, while the hind legs are in the stance, the middle legs are in swing. The front pair of legs are somewhat more irregular in timing and phase, as evident from the large variation depicted in the front leg data in Figure 2C. As suggested by Figure 1C, the insect’s abdomen is raised as its hind legs push against the ground, which, in side view, gives the impression that the mole cricket is bouncing backwards. Typically, the backward-bound gait is transiently demonstrated and the insect switches to double-tripod backward walking within up to 12 consecutive cycles of bound (6.0±2.5).

**Figure 2:**
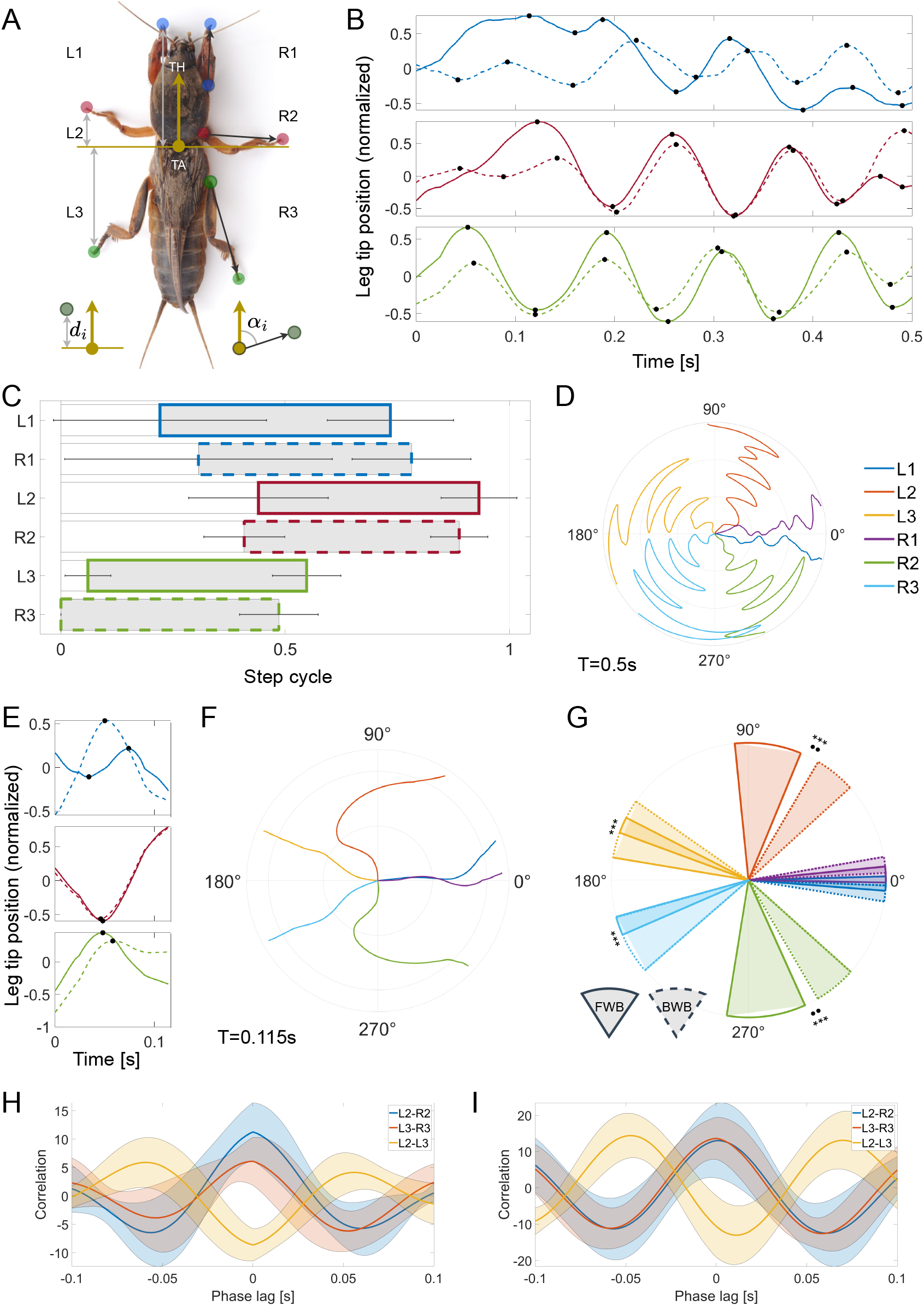
**A**. Parameters used in the mole cricket gait analysis. Colored dots depict base and tip of the three leg pairs. TA and TH: Thorax-Abdomen and Thorax-Head junctions, used to calculate leg position (*d*_*i*_; grey arrows, shown for left legs only) and leg angles (*a*_*i*_; black arrows, shown only for right legs). See Methods for details. **B**. An example of the leg-tip positions over time during backward-bound gait. Colors as in A. Solid lines – left legs; Dashed lines – right legs. **C**. Phasing of all legs relative to the right hind leg during backward bound. Colored bars show the average phase of the stance in normalized step cycle coordinates (extracted from the leg tips’ position data; see black dots in B), with standard deviation (N=20). **D**. An example of the leg angles during backward-bound (see A and Methods, same example as in B). Distance from the origin depicts time. **E**. An example of the leg-tip positions in time during forward-bound (ca. single cycle), *cf*. B. **F**. Leg angles during a cycle of forward-bound, *cf*. D. **G**. Legs’ angular coverage in forward vs. backward bound. Circular sectors show mean±STD of leg angles. Analysis based on data similar to that shown in D and F. Solid lines – forward-bound (N=10); Dotted lines – backward-bound (N, = 20). Differences in the mean angle, ••p<0.005; differences in the standard deviation, ***p<0.0005. **H-I**. Inter-leg phases (mean±STD) during the forward-and backward-bound gait (respectively) as demonstrated by cross-correlation of leg pairs.

Presenting a standing mole cricket with an aversive stimulus from behind, i.e., touching its rear parts with a similar decoy stimulus, sometimes resulted in a brief and transient (up to two cycles) forward-bound (Figs. 1D, 2E and F, N=10; 5 males and 5 females). A comparison of leg angles reveals significant differences in the trajectory of the middle and hind legs in forward vs. backward-bound (Fig. 2D, F, G): in forward-bound the middle legs span a larger angular sector and are oriented more perpendicularly to the body (a more posterior position), while the hind legs show altogether very little angular motion. The cross-correlation analysis presented in Figure 2H,I suggests overall similar phase relations to those described above for the backward-bound gait. Thus, despite similar temporal characteristics, the spatial analysis suggests that different legs take a dominant role in thrust generation in forward vs. backward-bound.

## Discussion

Mole crickets present an intriguing model system for the study of insect locomotion. Their cryptic life style in the subterranean environment greatly restricts our knowledge regarding their physiology and behavior. Their extreme morphological modifications, and specifically their leg adaptations, make any prediction regarding their modes of locomotion and walking gaits challenging. Based on the extreme and diverse adaptations seen in the different leg-pairs it is somewhat surprising that mole crickets predominately use the common double-tripod gait for walking (Zhang et al., 2011; 2019). However, it may be less surprising to have discovered that these fascinating insects present a unique, previously undescribed locomotion gait—the six-legged-bound gait.

Gait selection is a specific case of motor pattern selection (Pflüger, 2017; Pflüger et al., 2017). This concept is based on the accepted understanding that in vertebrates and invertebrates alike, most locomotion-related motor patterns are controlled by central nervous system neuronal networks known as central pattern generators (CPGs). CPG-generated rhythmic output induces an alternation between antagonistic motor neuron groups, resulting in motor behaviors with distinct phase relationships. Research in various invertebrate and vertebrate models has revealed common design principles of motor selection and decision-making: namely context-dependent selection of the appropriate motor network and the distinct network’s motor output (distinct phase scheme).

Hence, within the above framework, the bound gait reflects a specific functional architecture (functional connectivity) of the mole cricket’s locomotion CPGs, which is expressed in a distinct descending neuronal (brain) and neuro-modulatory context – in response to an aversive stimulus. Furthermore, our findings to date suggest that the bound gait constitutes an unstable state of the CPG networks that control locomotion in the insect (see Pflüger, 2017; Reches et al., 2019 for a discussion of network stability in insect locomotion). Future work will focus on the distinct conditions and mechanisms that enable the emergence of this fundamentally unique phase relations and coupling between the unit pattern-generator circuits controlling the different legs during bound (for a discussion of coupling in insect locomotion see David and Ayali, 2021; Serrano et al., 2024).

Animal escape responses are by definition fast and robust, comprising in principle a burst of muscle activation (Card, 2012; Branco and Redgrave, 2020). Widely-studied examples include the crayfish tail-flip behavior (e.g., Edwards, 1999; Krasne and Edwards, 2002) and the escape jump of the locust (Heitler and Burrows, 1977; Pearson and O’Shea, 1984). Escape responses, however, are not necessarily simple. They are the result of complex sensorimotor processes that include the processing of the input, the decision, the generation of a motor commands, and the activation of muscles that move the animal and maximizes its chances of survival (Domenici et al., 2011; Card, 2012). This is well exemplified by the much studied cockroach escape behavior (Camhi and Tom, 1978; Camhi et al., 1978). The herein described forward-bound bears some similarity to the evasive jump of the locust (as also seen in other orthopterans). However, in contrast to a jump that relies only on the hind legs (Heitler and Burrows, 1977), the forward-bound gait involves the alternation of at least the middle and hind pairs of legs, and is therefore very different and more complex. Though not common, mole crickets do jump or hop on occasion using their hind legs only. Similar to the case of the locust, in the adult mole cricket the jump will precede flight initiation (Amir and Ayali, unpublished). The backward bound, while clearly an evasive response, needs to be examined under a different framework, as it comprises a longer sequence of controlled leg movements demonstrating a unique pattern of coordination (compared to other locomotion gaits).

The novel bound gait described herein may be somewhat reminiscent of the previously reported galloping gait of the dung beetle (Smolka et al., 2013), as the galloping beetle also demonstrates a very unusual left-right legs’ synchrony. The case of the mole cricket is more conspicuous, however, for several major reasons, among them is the fact that the bound is one of several gaits available to the mole cricket, to be transiently selected at the appropriate context. This presents, of course, major challenges in control, as well as in biomechanics. Even more significant is the fact that unlike in the case of the beetle’s galloping (where the hind legs are dragged behind the insect without use), the bound gait involves all six legs of the insect, including the very dominant pair of hind legs.

The current study is part of a major endeavor aimed at constructing a detailed comparative description of the multiple locomotion gaits of the mole cricket. Such a comprehensive description is required in order to fully comprehend the six-legged-bound gait within the full scope of locomotion behaviors of this intriguing insect.

## Supporting information

Raw data used for all fgures

## Acknowledgments

We are very grateful to Alon Amir, Ron Jerbi, and Eran Rozen for their invaluable assistance in the early stages of this research project.

## Author contributions

Amir A. carried out the experiments and measurements. O.Y. processed the experimental data and performed the analysis. Ayali .A. conceived and supervised the project, wrote the manuscript with support from Amir A. and O.Y.

## Data availability

All raw data that served for generating the presented figures can be found in the supplementary file (Supplementary Table 1). Further data as required may be available upon request.

## Figures Legends

**Supplementary Table 1**

The table includes all data that served for generating the presented figures

**Supplementary Video S1**

The mole cricket’s backward-bound gait; speed x0.15

## References

Ayali, A., Borgmann, A., Büschges, A., Couzin-Fuchs, E., Daun-Gruhn, S., & Holmes, P. (2015). The comparative investigation of the stick insect and cockroach models in the study of insect locomotion. Current Opinion in Insect Science, 12, 1–10.

Biewener, A., & Patek, S. (2018). Animal locomotion. Oxford University Press.

Branco, T., & Redgrave, P. (2020). The neural basis of escape behavior in vertebrates. Annual review of neuroscience, 43(1), 417–439.

Camhi, J. M., & Tom, W. (1978). The escape behavior of the cockroach Periplaneta americana: I. Turning response to wind puffs. Journal of comparative physiology, 128, 193–201.

Camhi, J. M., Tom, W., & Volman, S. (1978). The escape behavior of the cockroach Periplaneta americana: II. Detection of natural predators by air displacement. Journal of comparative physiology, 128, 203–212.

Card, G. M. (2012). Escape behaviors in insects. Current opinion in neurobiology, 22(2), 180–186.

David, I., & Ayali, A. (2021). From Motor-Output to Connectivity: An In-Depth Study of in-vitro Rhythmic Patterns in the Cockroach Periplaneta americana. Frontiers in Insect Science, 1, 655933.

Domenici, P., Blagburn, J. M., & Bacon, J. P. (2011). Animal escapology I: theoretical issues and emerging trends in escape trajectories. Journal of Experimental Biology, 214(15), 2463–2473.

Edwards, D. H., Heitler, W. J., & Krasne, F. B. (1999). Fifty years of a command neuron: the neurobiology of escape behavior in the crayfish. Trends in neurosciences, 22(4), 153–161.

Graham, D. (1985). Pattern and control of walking in insects. In Advances in insect physiology (Vol. 18, pp. 31–140). Academic Press.

Heitler, W. J., & Burrows, M. (1977). The locust jump. I. The motor programme. Journal of Experimental Biology, 66(1), 203–219.

Irschick, D. J., & Garland Jr, T. (2001). Integrating function and ecology in studies of adaptation: investigations of locomotor capacity as a model system. Annual Review of Ecology and Systematics, 32(1), 367–396.

Kidd, J. (1825). X. On the anatomy of the mole-cricket. Philosophical transactions of the royal society of London, (115), 203–246.

Krasne, F. B., & Edwards, D. H. (2002). Crayfish escape behavior: lessons learned. In Crustacean experimental systems in neurobiology (pp. 3–22). Berlin, Heidelberg: Springer Berlin Heidelberg.

Mathis, A., Mamidanna, P., Cury, K. M., Abe, T., Murthy, V. N., Mathis, M. W., & Bethge, M. (2018). DeepLabCut: markerless pose estimation of user-defined body parts with deep learning. Nature neuroscience, 21(9), 1281–1289.

Pearson, K. G., & O’Shea, M. (1984). Escape behavior of the locust: the jump and its initiation by visual stimuli. In Neural mechanisms of startle behavior (pp. 163–178). Boston, MA: Springer US.

Pflüger, H. J. (2017). Motor pattern selection and initiation in invertebrates with an emphasis on insects. Neurobiology of Motor Control: Fundamental Concepts and New Directions, 195.

Pflüger, H. J., Grillner, S., & Robertson, B. (2017). Motor Pattern Selection. Neurobiology of Motor Control: Fundamental Concepts and New Directions, 177–223.

Reches, E., Knebel, D., Rillich, J., Ayali, A., & Barzel, B. (2019). The metastability of the double-tripod gait in locust locomotion. Iscience, 12, 53–65.

Ritzmann, R. E., & Büschges, A. (2007). Adaptive motor behavior in insects. Current opinion in neurobiology, 17(6), 629–636.

Smolka, J., Byrne, M. J., Scholtz, C. H., & Dacke, M. (2013). A new galloping gait in an insect. Current Biology, 23(20), R913–R915.

Serrano, S., Barrio, R., Lozano, Á., Mayora-Cebollero, A., & Vigara, R. (2024). Coupling of neurons favors the bursting behavior and the predominance of the tripod gait. Chaos, Solitons & Fractals, 184, 114928.

Wang, H., Lei, X., Zhang, G., Xu, C., Fang, Y., & Zhang, H. (2020). The earliest Gryllotalpinae (Insecta, Orthoptera, Gryllotalpidae) from mid-Cretaceous Burmese amber. Cretaceous Research, 107, 104292.

Xu, C., Fang, Y., & Wang, H. (2020). A new mole cricket (Orthoptera: Gryllotalpidae) from mid-Cretaceous Burmese amber. Cretaceous Research, 112, 104428.

Zhang, Y., Huang, H., Liu, X., & Ren, L. (2011). Kinematics of terrestrial locomotion in mole cricket gryllotalpa orientalis. Journal of Bionic Engineering, 8(2), 151–157.

Zhang, Y., Cao, J., Wang, Q., Wang, P., Zhu, Y., & Zhang, J. (2019). Motion characteristics of the appendages of mole crickets during burrowing. Journal of Bionic Engineering, 16, 319–327.

