## Supplementary material for "Six-legged-bound: a newly-described insect gait": Raw data used for all fgures

forward_bound

| | L1 | R1 | L2 | R2 | L3 | R3 |
|---|---|---|---|---|---|---|
| Leg angle - Mean | 0.138700766 | -0.315104471 | 1.697966635 | -1.626847764 | 2.830471781 | -2.668863517 |
| | 0.002803463 | -0.077189977 | 1.368793297 | -1.436778479 | 2.706081475 | -2.610238561 |
| | 0.13280201 | 0.062543125 | 1.760696863 | -1.505357607 | 2.785390381 | -2.654839794 |
| | -0.221114045 | 0.169678578 | 1.202847761 | -1.735175104 | 2.960614068 | -2.941759894 |
| | -0.123933251 | 0.005337391 | 1.346358499 | -1.355334221 | 2.641072971 | -2.794876006 |
| | -0.027958675 | 0.257064096 | 0.946525865 | -1.010128114 | 2.645041486 | -2.990036833 |
| | -0.135967034 | 0.117388491 | 1.692934681 | -1.587573372 | 2.749609304 | -2.99155106 |
| | 0.08819997 | 0.077893079 | 1.417846634 | -1.202209106 | 2.521164438 | -2.727312598 |
| | 0.002023853 | 0.017334477 | 1.471968013 | -1.408955886 | 2.578620732 | -2.792407548 |
| | -0.110851626 | 0.087522614 | 1.342432606 | -1.509313104 | 2.874119358 | -2.845772681 |
| Leg angle - standard deviation | 0.237889839 | 0.219024792 | 0.506450557 | 0.497747989 | 0.073165962 | 0.144521413 |
| | 0.046917034 | 0.085307208 | 0.420143336 | 0.672876376 | 0.176484508 | 0.125821367 |
| | 0.11080011 | 0.103034471 | 0.53471912 | 0.532891015 | 0.128191266 | 0.117105735 |
| | 0.041972301 | 0.058802057 | 0.476786961 | 0.561244603 | 0.113668426 | 0.086499739 |
| | 0.110645567 | 0.126461528 | 0.391313927 | 0.377711909 | 0.127322001 | 0.085428789 |
| | 0.059581477 | 0.134996812 | 0.398903247 | 0.484759187 | 0.121808764 | 0.165560462 |
| | 0.097159918 | 0.084450322 | 0.672245632 | 0.619403814 | 0.076226828 | 0.184404903 |
| | 0.09636112 | 0.066554369 | 0.368170543 | 0.684906351 | 0.12523007 | 0.101776852 |
| | 0.145770844 | 0.165702958 | 0.415117792 | 0.711777748 | 0.153045887 | 0.131610781 |
| | 0.100826127 | 0.126936845 | 0.72385932 | 0.692815619 | 0.094322687 | 0.249787583 |

backward_bound

| | L1 | R1 | L2 | R2 | L3 | R3 |
|---|---|---|---|---|---|---|
| Leg angle - Mean | -0.171138845 | 0.1621079 | 0.95583316 | -0.922989389 | 2.644343337 | -2.455087188 |
| | -0.107632616 | -0.03624846 | 0.691932796 | -1.25757179 | 2.643298828 | -2.904535768 |
| | -0.100205274 | 0.078660898 | 1.295516833 | -0.756540161 | 2.823722334 | -2.570176168 |
| | -0.146077919 | 0.037855713 | 0.742880098 | -0.92845729 | 2.877522764 | -2.710977813 |
| | -0.14193235 | 0.098225812 | 0.761408633 | -1.217406934 | 2.734628781 | -2.648467699 |
| | -0.153338092 | -0.084992058 | 0.7280622 | -0.994960519 | 2.778775728 | -2.788660026 |
| | -0.147462329 | 0.269736647 | 0.935249982 | -0.895958834 | 2.583168289 | -2.693762695 |
| | -0.103475919 | 0.137689389 | 0.826263675 | -0.941442861 | 2.796725456 | -2.91419393 |
| | 0.250191389 | 0.309945628 | 1.297856071 | -0.723111955 | -3.036560589 | -2.293363036 |
| | -0.068918255 | -0.03502228 | 0.661446472 | -1.048176126 | 2.92920112 | -2.631244285 |
| | -0.030033205 | 0.057257047 | 0.904278617 | -0.896252725 | 2.776380188 | -2.663246088 |
| | 0.05897452 | 0.211204196 | 1.121423908 | -0.609875929 | 2.725763321 | -2.302677712 |
| | 0.023974773 | 0.111704887 | 0.994989665 | -0.894676181 | 2.83297644 | -2.646374079 |
| | -0.238728129 | 0.061770397 | 0.944566653 | -0.717637208 | 2.723083499 | -2.601977445 |
| | -0.135315296 | 0.155649963 | 0.817100834 | -0.951116455 | 2.654203448 | -2.854271887 |
| | -0.175538681 | 0.029670933 | 0.661767325 | -0.955900338 | 2.582878931 | -2.596148416 |
| | -0.158563014 | -0.07115549 | 0.846390657 | -0.730512508 | 2.678734443 | -2.811901219 |
| | -0.08877399 | 0.22129648 | 1.103789314 | -0.591977197 | 2.841421668 | -2.315953073 |
| | -0.1625538 | 0.270906323 | 0.802279313 | -0.855543961 | 2.538043428 | -2.725114044 |
| | -0.147866212 | 0.239884175 | 0.692815715 | -0.840377157 | 2.676124225 | -2.770367329 |
| Leg angle - standard deviation | 0.09482172 | 0.139408612 | 0.336453593 | 0.339048455 | 0.567116677 | 0.584522821 |
| | 0.141145344 | 0.102888425 | 0.312428806 | 0.2441258 | 0.350778392 | 0.375999795 |
| | 0.104039597 | 0.104897923 | 0.28740014 | 0.419039813 | 0.538306738 | 0.511513625 |
| | 0.104734911 | 0.154198021 | 0.290671158 | 0.174555061 | 0.37679482 | 0.344862721 |
| | 0.091091776 | 0.11726811 | 0.179616865 | 0.395510679 | 0.217406166 | 0.489851375 |
| | 0.083071863 | 0.083493984 | 0.322035286 | 0.164344507 | 0.243798387 | 0.475833201 |
| | 0.079614127 | 0.091061416 | 0.320609438 | 0.278607942 | 0.660272769 | 0.366325713 |
| | 0.060841333 | 0.098985696 | 0.262154878 | 0.16705532 | 0.357031318 | 0.253402503 |
| | 0.249766526 | 0.132604895 | 0.37581437 | 0.523338551 | 0.438176008 | 0.602888913 |
| | 0.098740514 | 0.091201054 | 0.227123579 | 0.304744321 | 0.309088521 | 0.35559294 |
| | 0.097544023 | 0.118300644 | 0.416656806 | 0.30781004 | 0.315202136 | 0.369687134 |
| | 0.162076262 | 0.155353735 | 0.200319004 | 0.402361834 | 0.488318343 | 0.438845757 |
| | 0.082912975 | 0.111172369 | 0.285047209 | 0.306985587 | 0.358335829 | 0.360106573 |
| | 0.146718845 | 0.092469129 | 0.352885774 | 0.276244331 | 0.532161422 | 0.497160228 |
| | 0.091931814 | 0.207428222 | 0.236039595 | 0.275685191 | 0.292985464 | 0.319379595 |
| | 0.13942943 | 0.162614092 | 0.39604442 | 0.390028493 | 0.736172118 | 0.799886965 |
| | 0.174117164 | 0.04647239 | 0.287018295 | 0.24090247 | 0.386883216 | 0.269944531 |
| | 0.188981995 | 0.133502511 | 0.259714722 | 0.406445881 | 0.754113833 | 0.48859752 |
| | 0.08263485 | 0.103609988 | 0.362859683 | 0.302561897 | 0.527899175 | 0.430809912 |
| | 0.081567478 | 0.110377807 | 0.270592526 | 0.295210446 | 0.469893499 | 0.493709617 |
